# Quantification of Monosynaptic Rabies Tracing Efficiency

**DOI:** 10.1101/2022.08.31.506012

**Authors:** Maribel Patiño, Willian N. Lagos, Neelakshi S. Patne, Paula A. Miyazaki, Edward M. Callaway

## Abstract

Retrograde monosynaptic tracing using glycoprotein-deleted rabies virus is an important component of the toolkit for investigation of neural circuit structure and connectivity. It allows for the identification of first-order presynaptic connections to cell populations of interest across both the central and peripheral nervous system, helping to decipher the complex connectivity patterns of neural networks that give rise to brain function. Despite its utility, the efficiency with which genetically modified rabies virus spreads retrogradely across synapses remains uncertain. While past studies have revealed conditions that can increase or decrease the numbers of presynaptic cells labeled, it is unknown what proportion of total inputs to a starter cell of interest are labeled. It is also unknown whether synapses that are more proximal or distal to the cell body are labeled with different efficiencies. Here we use a new rabies virus construct that allows for the simultaneous labeling of pre and postsynaptic specializations to quantify efficiency of spread at the synaptic level in mouse primary visual cortex. We demonstrate that with typical conditions about 40% of first-order presynaptic excitatory inputs are labeled. We show that using matched tracing conditions there is similar efficiency of spread from excitatory or inhibitory starter cell types. Furthermore, we find no difference in the efficiency of labeling of excitatory inputs to postsynaptic sites at different subcellular locations.

## INTRODUCTION

Monosynaptic rabies tracing using glycoprotein (G) – deleted rabies virus (RVdG) is a powerful tool for the study of neural circuit connectivity. This method enables scientists to label, genetically manipulate, or monitor the activity of brain-wide monosynaptic inputs to cell populations of interest. Since its introduction (Wickersham, Finke, et al., 2007; Wickersham, Lyon, et al., 2007) it has been widely used for the identification of presynaptic inputs to single neurons (Marshel et al., 2010; Rancz et al., 2011; Rossi et al., 2020; Wertz et al., 2015), projection-defined neurons (Cruz-Martín et al., 2014; Levine et al., 2014), adult-born neurons (Deshpande et al., 2013; Garcia et al., 2014), transplanted neurons (Doerr et al., 2017; Grealish et al., 2015), hPSC-derived organoid neurons (Andersen et al., 2020; Miura et al., 2020), and genetically defined excitatory neurons (DeNardo et al., 2015; Kim et al., 2015), inhibitory neurons (Miyamichi et al., 2013; Wall et al., 2016), and non-neuronal cell types (Clark et al., 2021; Mount et al., 2019). In addition to being used for the identification of inputs, the incorporation of Ca^2+^ indicators and light-activated opsins into rabies reagents (Osakada et al., 2011) allows rabies tracing experiments to probe the relationship between function and connectivity (Rossi et al., 2020; Tian et al., 2016; Wertz et al., 2015; Wester et al., 2019). The ability of genetically modified RVdG to selectively spread retrogradely between synaptically connected cells allows for the identification of presynaptic partners regardless of their distance from one another and has led to novel insights throughout the nervous system.

Despite the utility and widespread use of monosynaptic rabies tracing to study neural connectivity, there is uncertainty about the efficiency of transsynaptic spread from starter cells to input neurons. Although studies quantifying inputs to single neurons have provided some insight into the efficiency of spread (Marshel et al., 2010; Miyamichi et al., 2011; Rancz et al., 2011), results vary widely across experimental conditions and no direct measurements of spread efficiency are available. Furthermore, using the convergence index, defined as the number of rabies-labeled input neurons divided by the number of starter cells, recent studies have shown that the number of input neurons labeled per starter cell can be improved about 10-fold by modifying the rabies glycoprotein (Kim et al., 2016) or replacing the rabies virus strain (Reardon et al., 2016). Despite allowing quantitative comparisons between different reagents and conditions, these studies relied on methods that result in large animal to animal variability, require tedious counting across many animals, and can display high variability depending on the number of starter neurons (Tran-Van-Minh et al., 2022). Most importantly, convergence index measurements fail to quantify what proportion of all inputs are labeled. Additionally, the use of convergence index fails to address questions of the influence of biological factors on rabies spread efficiency, such as distance of synapses to the starter cell soma or differences of spread from different starter cell types.

In this study we examine the efficiency of RVdG retrograde spread from starter cell to input neurons at the synaptic level. We designed a new genetically modified rabies virus that labels presynaptic and excitatory postsynaptic densities. This construct allows us to quantify the proportion of excitatory synapses on a starter cell that have their corresponding input neuron labeled with rabies virus. We find that with the particular reagents and conditions that we used, rabies retrogradely labels about 35% to 40% of excitatory inputs to each of the excitatory and inhibitory cell populations we tested. Furthermore, we found that efficiency of spread does not vary by the proximity or distance of synapses to the starter cell soma or by the type of neuronal dendrite. Overall, this study provides insight into some long-standing questions about the efficiency of monosynaptic rabies tracing. We further discuss that desired tracing efficiency varies depending on experimental aims and that efficiency was not maximized with the experimental design used here.

## RESULTS

### RVdG-PSD95GFP-SynPhRFP viral construct allows simultaneous fluorescent labeling of pre and postsynaptic specializations

To investigate the efficiency of rabies transsynaptic spread from starter cells to input neurons we developed a high throughput and precise method to directly measure transsynaptic spread at the synaptic level. We created a new deletion-mutant rabies virus construct that expresses two synaptic fusion protein transgenes from the rabies G locus (Figure 1A), modeled after existing rabies constructs known to label multiple subcellular compartments of neurons (Wickersham et al., 2013). One gene encodes for a fusion protein of the excitatory postsynaptic marker, postsynaptic density 95 (PSD-95) and enhanced green fluorescent protein (eGFP). The other encodes for a fusion protein of the presynaptic marker synaptophysin (SynPh) and the bright and photostable red fluorescent protein TagRFP-T (Shaner et al., 2008). We chose the synaptophysin-TagRFP-T (SynPhRFP) fusion protein as it has previously been shown to result in a punctate red fluorescent pattern that colocalizes with varicosities on axons when expressed from the rabies genome (Wickersham et al., 2013). Injection of EnvA+ RVdG-PSD95GFP-SynPhRFP in primary visual cortex (V1) of Sim1-Cre mice expressing TVA and oG in Cre^+^ layer 5 neurons (derived from AAV helper viruses, see Methods) resulted in strong punctate green and red fluorescent labeling (Figure 1B), with green labeling prominent on dendrites and dendritic spines (Figure 2E, 2F). Super-resolution Airyscan imaging (see methods) revealed that GFP labeling colocalized with a subset of the puncta labeled with antibody staining against endogenous PSD-95 (Figure 1C). In addition to synaptic labeling there were typically nuclear aggregates of both red and green fluorescent protein in infected neurons (Figure 1B).

**Figure 1:**
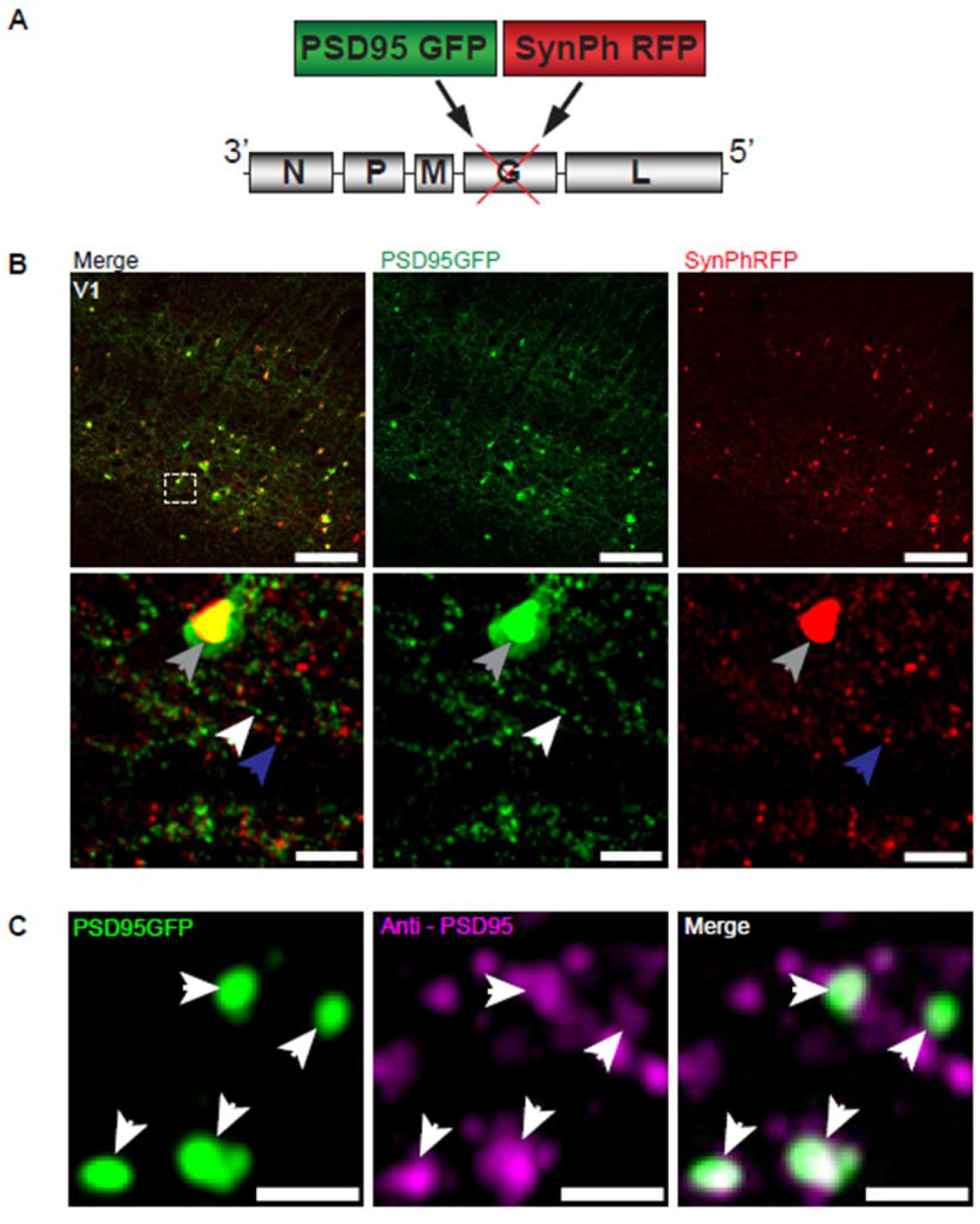
RVdG-PSD95GFP-SynPhRFP allows simultaneous fluorescent labeling of pre and postsynaptic densities. (A) Schematic of RVdG-PSD95GFP-SynPhRFP viral construct design. Two transgenes were inserted into the G locus of the rabies genome. One encodes a presynaptically-targeted fluorescent fusion protein, synaptophysin TagRFP-T (SynPhRFP), and the other a postsynaptically targeted fluorescent fusion protein, PSD-95 eGFP (PSD95GFP). (B) Coronal sections of Sim1-Cre mouse expressing TVA and oG in V1 infected with EnvA+ RVdG-PSD95GFP-SynPhRFP imaged at 20X with confocal microscopy. Top row shows neurons expressing both PSD95GFP and SynPhRFP fusion proteins. Bottom row shows a zoomed in image of the region enclosed by the dashed square in the top row. White arrows point to PSD-95 puncta, blue arrows to synaptophysin puncta, and gray arrows to large non-specific nuclear fluorescent aggregates. Scale bars represent 100 um (top row) or 10 um (bottom row). (C) Airyscan super-resolution images taken at 63X showing co-localization of PSD95GFP fusion protein expressed from the rabies genome with anti-PSD95 antibody staining in magenta. Scale bar = 1 um.

**Figure 2:**
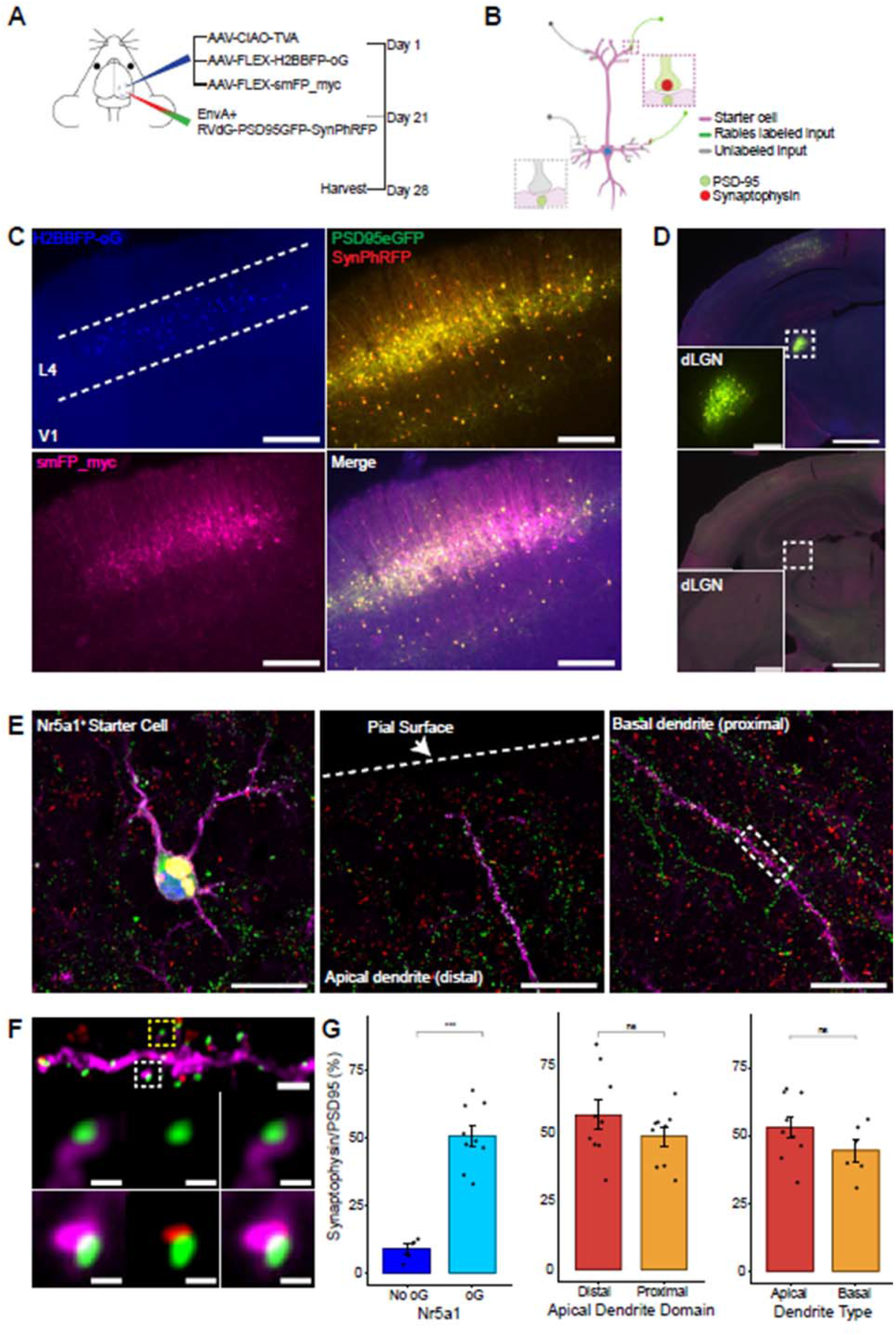
Efficiency of transsynaptic spread from excitatory L4 Nr5a1+ starter cells to excitatory inputs. (A) Schematic illustration of experimental design and timeline for monosynaptic rabies tracing. Nr5a1-Cre mice were injected in V1 with a mixture of AAV-CIAO-TVA, AAV-FLEX-H2BBFP-oG, and AAV-FLEX-smFP_myc. 3 weeks later EnvA+ RVdG-PSD95GFP-SynPhRFP was injected into the same site and allowed to express for 7 days. (B) Schematic of rabies retrograde spread efficiency quantification paradigm. Starter neurons are distinguished from input neurons based on expression of nuclear BFP from AAV-FLEX-H2BBFP-oG in addition to fusion proteins from EnvA+ RVdG-PSD95GFP-SynPhRFP. Starter neurons expressing smFP_myc are used for synaptic quantification to allow tracing of distal dendrites. Retrograde spread efficiency is measured by quantifying the proportion of postsynaptic densities (PSD95GFP) on the starter neuron opposed with rabies-labeled presynaptic terminals (SynPhRFP). (C) Representative example images of V1 injection site, obtained using widefield fluorescence microscopy at 10X magnification. Scale bar = 200um. (D) Coronal section example images obtained using widefield fluorescence microscopy at 10X showing long range monosynaptic input neurons in dLGN to Nr5a1^+^ L4 neurons in V1 when using the new RVdG construct (top). No retrograde spread is observed when glycoprotein is omitted, see Supplementary Figure 1 for additional information. Insets are zoomed in images of dashed box regions. Scale bar represents 1 mm in hemisection image or 200 um in inset. (E) Images obtained using Airyscan super-resolution imaging at 63X. Left, example image of starter neuron (H2BBFP+,PSD95GFP+, and SynPhRFP+) labeled with smFP_myc. Middle, example image of the distal domain of an apical dendrite of a starter neuron. Right, example image of the proximal domain of a basal dendrite. Scale bar = 20um (all three). (F) Spatial resolution using Airyscan imaging is sufficient to quantify rabies transsynaptic spread at the synapse level. Zoomed in image of boxed region in (E) right, illustrating PSD-95 puncta co-colocalized with cytoplasmic smFP_myc. Top, yellow boxed region highlights a spine with PSD-95 puncta without an opposed rabies-labeled presynaptic density. White boxed region highlights a spine with PSD-95 puncta with an opposed rabies-labeled presynaptic density. Middle row, zoomed in images of yellow boxed region and bottom row are zoomed in images of white boxed region. Top, scale bar = 2um and middle and bottom scale bar = 0.5 um. (G) Percent of postsynaptic densities (PSD95GFP) on Nr5a1^+^ starter cells opposed with rabies-labeled presynaptic terminals (SynPhRFP). Left, quantification of colocalization at baseline (no glycoprotein) due to L4 to L4 connections compared to colocalization from transsynaptic spread (with glycoprotein). Middle, colocalization on the distal vs proximal domains of apical dendrites. Right, colocalization on apical vs basal dendrites. Values are reported as mean ± SEM. Statistics were calculated from the Wilcoxon rank-sum test for non-parametric comparisons. Individual data points (circles) indicate values for each neuron. ns, not significant.

### Rabies transsynaptic spread efficiency across excitatory synapses from excitatory starter neurons

To quantify the proportion of inputs labeled with rabies virus we injected a mixture of three Cre-dependent helper adeno-associated viruses (AAV) into the primary visual cortex (V1) of Nr5a1-Cre mice to target initial rabies infection to a sparse population of L4 excitatory neurons (Harris et al., 2014) (Figure 2A, 2B). 1) AAV8-nef-AO-66/71-TVA950 (AAV-CIAO-TVA), expresses the TVA receptor for the avian sarcoma leukosis virus envelope protein, EnvA, which is necessary for entry of pseudotyped EnvA^+^ RVdG into Cre^+^ cells. However, recombinant-independent off-target leak expression of transgenes is common in recombinase-dependent DIO and FLEX AAV constructs. Because even miniscule quantities of TVA leak expression is sufficient for pseudotyped rabies to enter off-target cells and confound results (Callaway & Luo, 2015), it was necessary to minimize leak expression of TVA. We therefore used the novel cross-over insensitive ATG-out (CIAO) AAV construct, which has been shown to nearly eliminate leak expression and provide reliable and targeted transgene expression (Fischer et al., 2019), to express TVA in our population of interest. Experiments in which AAV-CIAO-TVA and EnvA+ RVdG were injected into V1 of Cre-negative mice resulted in 15.83 ± 11.33 rabies-infected neurons (mean ± SEM, n = 3 mice) indicating low levels of Cre-independent TVA expression. 2) AAV8-hSyn-FLEX-H2BBFP-oG expresses optimized rabies glycoprotein (oG) (Kim et al., 2016), which allows for trans-complementation in EnvA+ RVdG infected neurons, also termed starter cells, allowing the rabies to spread retrogradely into presynaptically connected inputs. Co-expression of nuclear mTagBFP2 (BFP), a brighter and more photostable blue fluorescent protein (Subach et al., 2008), and oG allows for the unambiguous identification of starter cells (Figure 2C, 2E), defined as any cell that expresses both oG and rabies transgenes. 3) Because H2BBFP only labels the nucleus of starter cells, we also used AAV-CAG-FLEX-smFP_myc to generate cytoplasmic labeling, allowing dendrites of starter cells to be traced. This AAV expresses spaghetti monster fluorescent protein (smFP), which consists of dark non-fluorescent GFP fused to 10 Myc epitope tags that can be combined with anti-Myc antibody staining to attain bright fluorescent labeling of subcellular structures (Viswanathan et al., 2015). Infection of L4 excitatory neurons with AAV-CAG-FLEX-smFP_myc resulted in strong dendritic labeling that improved the ability to accurately trace apical and basal dendrites of starter cells (Figure 2C, 2E, 2F). Three weeks after injection of a mixture of the three AAV helper viruses, EnvA^+^ RVdG-PSD95GFP-SynPhRFP was injected into the same location in V1 and allowed to express for 7 days. As expected from transsynaptic spread, we observed that many thalamic dorsal lateral geniculate nucleus (dLGN) long-distance inputs to L4 excitatory neurons were reliably labeled with RVdG-PSD95GFP-SynPhRFP (Figure 2D).

To quantify the proportion of inputs labeled on starter neurons we first identified starter neurons that also expressed smFP using wide-field fluorescence microscopy. Select areas of starter neuron dendrites that could be traced back to their parent cell bodies were imaged using Airyscan super-resolution microscopy (Huff et al., 2017) to increase the resolution and signal-to-noise and allow visualization of single synaptic puncta (Figure 2F). To assess the transsynaptic spreading efficiency, we quantified the proportion of starter cell excitatory postsynaptic specializations, labeled with PSD95GFP, that were directly opposed with rabies-labeled presynaptic terminals, labeled with SynPhRFP (Figure 2F). Because L4 excitatory neurons connect to one another, the direct connections between starter cells can generate co-labeling of pre- and postsynaptic specializations independent from transsynaptic spread. It was therefore important to begin by quantifying how much of this “background label” is present under conditions in which similar numbers of L4 neurons are directly infected with EnvA^+^ RVdG-PSD95GFP-SynPhRFP, but there is no transsynaptic spread. To quantify this we calculated the proportion of PSD95GFP puncta opposed to SynPhRFP in experiments that omitted oG (no AAV8-hSyn-FLEX-H2BBFP-oG, Supplementary Figure 1A, 1B, 1C). The omission of oG prevents retrograde spread of rabies virus and labeling of input neurons (Figure 2D), therefore observed SynPhRFP colocalization with PSD95GFP must be a result of L4 to L4 excitatory starter neuron connections. We found that neurons in which oG was used for trans-complementation displayed significantly higher proportions of PSD95GFP puncta colocalized with SynPhRFP compared to neurons in which oG was omitted (50.61 ± 3.92% versus 8.86 ± 1.79% mean ± SEM respectively, Wilcoxon rank-sum test, p = 0.001, n = 9 neurons across 3 mice and n = 5 neurons across 2 mice; Figure 2G). Within experimental condition groups, puncta colocalization did not vary significantly across neurons from distinct experimental animals. The difference between these values (50.61% - 8.86%) yields an estimate of 42% of postsynaptic densities whose presynaptic partners are transsynaptically labeled.

To determine how efficiency of spread is related to distance from the cell body we sampled from different portions of the dendritic arbors of L4 starter cells. Sampled areas within 75um of the soma were classified as proximal dendritic domains and areas within 75um of the pial surface were considered distal domains. These distal domains are estimated to be about 250 to 350um from the soma. We found no significant differences between efficiency of spread at proximal regions of apical dendrites compared to the distal regions as determined by PSD95GFP and SynPhRFP puncta colocalization (49.14 ± 3.49% versus 57 ± 5.43% respectively, Wilcoxon rank-sum test, p = 0.34; n = 9 neurons across 3 mice, Figure 2G). We also compared labeling efficiency at basal dendrites versus apical dendrites and observed no significant difference (44.52 ± 3.96% versus 53.05 ± 3.99% respectively, Wilcoxon rank-sum test, p = 0.22; n = 9 neurons across 3 mice and n = 6 neurons across 3 mice, Figure 2G).

**Supplementary Figure 1:**
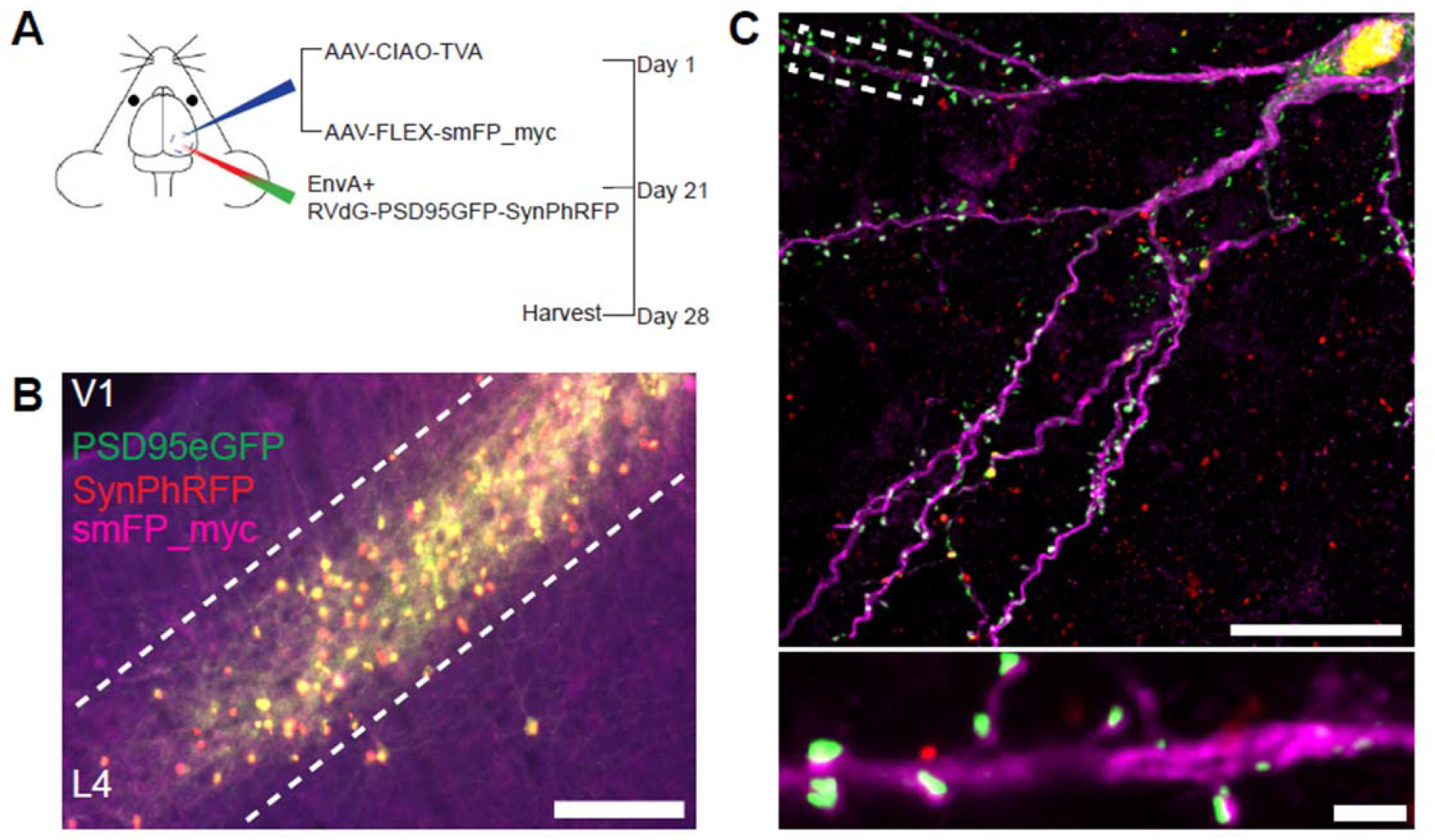
Experimental design for control experiments omitting oG, related to figure 2. (A) Schematic illustration of experimental design and timeline for control experiments without glycoprotein. Nr5a1-Cre mice were injected in V1 with a mixture of AAV-CIAO-TVA and AAV-FLEX-smFP-myc. 3 weeks later EnvA+ RVdG-PSD95GFP-SynPhRFP was injected into the same site and allowed to express for 7 days. (B) Representative image of V1 injection site, obtained using widefield fluorescence microscopy at 10X. Scale bar = 200um. (C) Images obtained using Airyscan super-resolution imaging at 63X. Top, example image of rabies-infected neuron labeled with smFP_myc, without glycoprotein. Bottom, zoomed in image of boxed region in top image, illustrating PSD-95 puncta colocalized with cytoplasmic smFP_myc. Scale bar = 20um (top) and scale bar = 2um (bottom).

### Rabies transsynaptic efficiency across excitatory synapses from inhibitory starter neurons

To assess possible differences in efficiency of rabies retrograde spread for different starter cell types, we conducted monosynaptic rabies tracing using RVdG-PSD95GFP-SynPhRFP as described above, but using mouse lines that express Cre recombinase in two distinct classes of inhibitory neurons. We used the knock-in Som-IRES-Cre and Vip-IRES-Cre lines to target initial infection to either somatostatin (Sst)-expressing or vasoactive intestinal peptide (Vip)-expressing inhibitory neurons (Figure 3A). We selected Sst and Vip interneurons as both exhibit very low levels of recurrent connections compared to parvalbumin (Pvalb)- expressing neurons, which connect extensively to each other (Campagnola et al., 2022; Pfeffer et al., 2013). We found no significant difference between the percent of PSD95GFP puncta apposed to SynPhRFP on Sst dendrites compared to Vip dendrites (36.13 ± 1.73% versus 38.38± 1.22% respectively, Wilcoxon rank-sum test, p = 0.44, n = 9 neurons across 3 mice per line; Figure 3C). These values are comparable to the efficiency of labeling inputs to L4 starter cells (42%, see above). We also assessed whether retrograde spread efficiency varied based on distance from the cell body. For Sst starter cells, colocalization of PSD95GFP with SynPhRFP on proximal dendritic domains (within 75um of the cell body) did not differ significantly from distal dendritic domains (within 75um of the pial surface) (37.50 ± 1.93% versus 33.85 ± 2.43% respectively, Wilcoxon rank-sum test, p = 0.55). We observed similar results for Vip starter cells, with no difference between retrograde spread efficiency at proximal versus distal domains (39.80 ± 1.90% versus 36.122 ± 1.14% respectively, Wilcoxon rank-sum test, p = 0.11)

**Figure 3:**
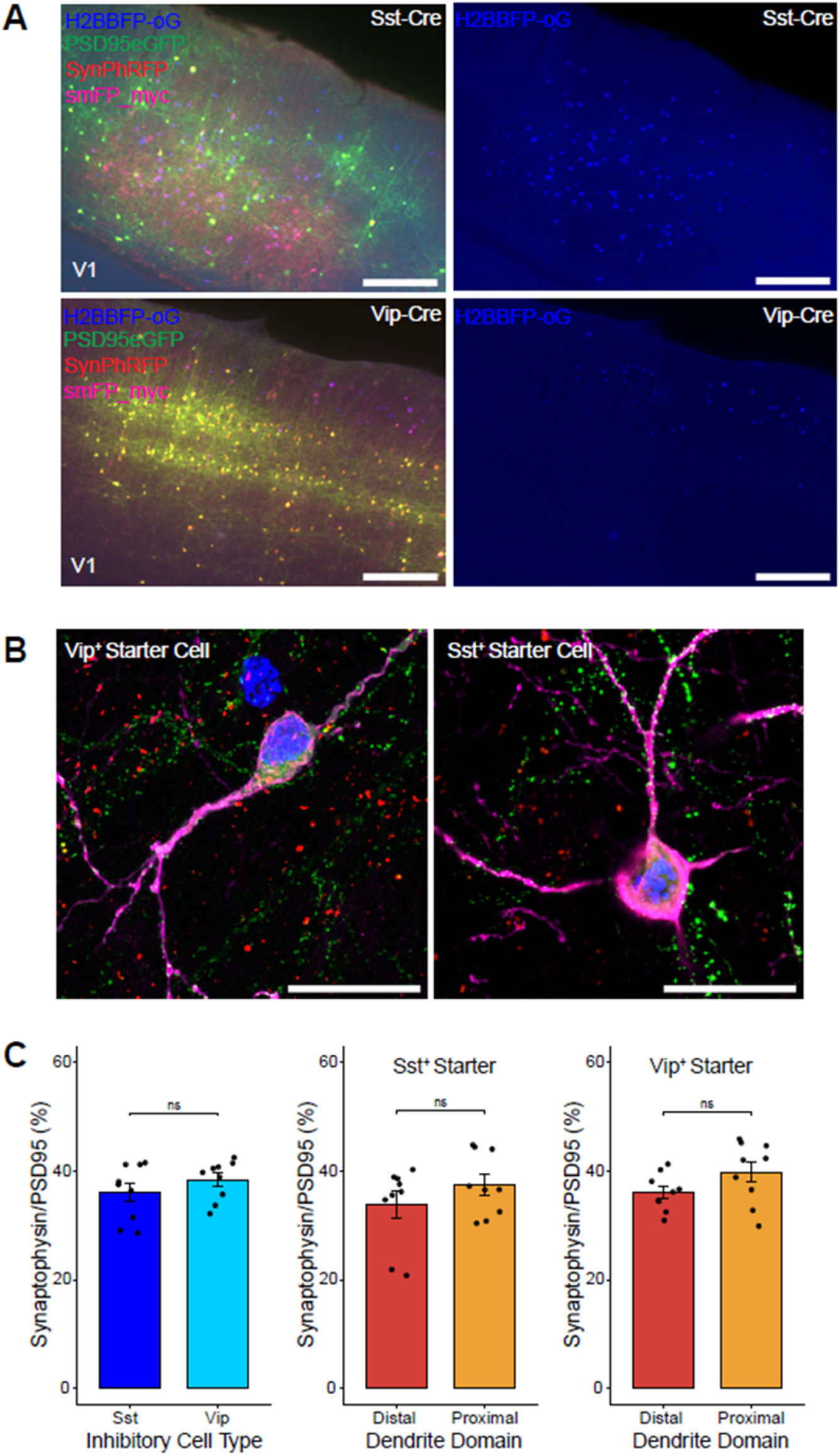
Efficiency of transsynaptic spread from inhibitory starter cells to excitatory inputs. (A) Representative example images of V1 injection site for Sst-Cre mouse line (top) and Vip-Cre mouse line (bottom), obtained using widefield fluorescence microscopy at 10X. Scale bar = 200um. (B) Images obtained using Airyscan super-resolution imaging at 63X. Example image of Vip+ (right) and Sst+ (left) starter neurons (H2BBFP+,PSD95GFP+, and SynPhRFP+) labeled with smFP_myc. Scale bar = 20um. (C) Left, percent of postsynaptic densities (PSD95GFP) on VIP+ and Sst+ starter cells opposed with rabies-labeled presynaptic terminals (SynPhRFP). Middle, colocalization on the distal vs proximal domains of Sst+ dendrites. Right, colocalization on the distal vs proximal domains of Vip+ dendrites. Values are reported as mean ± SEM. Statistics were calculated from Wilcoxon rank-sum test for non-parametric comparisons. Individual data points (circles) indicate values for each neuron. ns, not significant.

## DISCUSSION

Genetically modified RVdG has proven instrumental in deciphering the intricate connectivity patterns of neural networks. Here we investigated the efficiency with which RVdG spreads retrogradely across excitatory synapses and report, for the first time, direct measurements of the proportion of starter neurons’ postsynaptic densities whose presynaptic inputs are labeled using the monosynaptic rabies tracing system. Our results demonstrate that with the particular AAV helper viruses, AAV titers, and EnvA^+^RVdG titers and strain we used, about 35% to 40% of excitatory inputs to the examined cell types were labeled.

Of particular note, we found that efficiency of spread did not vary depending on the distance of synapses to the cell body or across dendrite types. We were concerned that more distal synapses might be less likely to be labeled than proximal synapses because antibody staining for rabies particles shows that during the first 1-3 days post-infection, particles are concentrated around the cell body; but they are not detected more distally until later time points (Nassi & Callaway, 2006). In the experiments described here, efficiency of spread was quantified at 7 days post-infection. It therefore appears that any initial bias for spread at proximal synapses might be eliminated by use of longer survival times. Since we only compared efficiency at one typical time point, care should be taken in interpreting results at shorter times post-infection.

Previous studies quantifying the numbers of rabies labeled input cells per starter cell (convergence index) have identified several different factors that can increase or decrease the efficiency of transsynaptic spread. While there are few direct comparisons with matched titers of rabies viruses and helper viruses, both the strain of rabies virus and the strain and variant of rabies glycoprotein are likely to be influential. Importantly, when a particular rabies strain and glycoprotein are used, the convergence index appears to depend most strongly on the levels of G expressed in the starter cells (Callaway & Luo, 2015). Studies suggest that maximizing G expression, either through the use of strong promoters (Miyamichi et al., 2013) or optimized helper virus concentration (Lavin et al., 2020), increases spread efficiency. Importantly, the use of 2A linker elements has been implicated to reduce expression levels of G (Wall et al., 2010; Watabe-Uchida et al., 2012). Thus, the use of separate AAVs, with one expressing glycoprotein independently from other genes, is recommended to improve efficiency.

A second factor likely to influence efficiency is the number of rabies particles entering a starter cell (Callaway & Luo, 2015). The variability between individual cells that we sampled (Figure 2G and 3C) is likely to reflect these differences. For example, there is expected variability in the numbers of oG-expressing AAV copies and EnvA^+^ RVdG particles entering each starter cell. Despite the variability that we observe between individual cells, under matched conditions these differences converge to similar means when averaged across even modest numbers of cells (typically 9 cells in our sample) that are far lower than the number of total starter cells in a typical animal. It is noteworthy that it would be straightforward to increase the efficiency well above the 35-40% we observed by simply increasing the levels of oG expression. For example, our AAV helper viruses were designed to allow unambiguous detection of starter cells for accurate quantification, and we therefore utilized a 2A linker element to express both nuclear BFP and oG from a single helper virus. For experiments where unambiguous starter cell detection is not crucial, oG expression can be increased by expression as a single protein from an alternate helper virus. Additionally, oG expression can be further increased through use of an expression amplification system such as TRE/tTA (Lavin et al., 2020).

These observations suggest that the efficiency of rabies tracing can be manipulated over a wide range depending on experimental goals. But counterintuitively, the ideal rate of spread is probably far less than 100% for most experiments. This is because the spread of rabies is likely to depend on the numbers of synaptic contacts as well as the size of the postsynaptic density, both of which are related to functional connection strength (Holler et al., 2021; Murthy et al., 2001). Thus, if a reasonably small fraction of inputs were labeled, the numbers of labeled presynaptic cells of different types would be expected to scale with the strength of connections from each cell type. But if rabies were to spread across 100% of synaptic contacts, this sensitivity would be lost; every input cell would be labeled regardless of connection strength. On the other hand, if efficiency were exceedingly low one would expect more variability between animals and the potential to fail to detect inputs from weak connections. We suggest that values in the range of 35-50% of synaptic contacts labeled are likely to be near optimal in providing both large numbers of labeled input neurons and some degree of sensitivity to connection strength (Henrich et al., 2020).

Considering the finding that rabies labels less than half of synaptic inputs, it is likely that differential spread of rabies virus leads to differences in the probability of distinct input types being labeled. Although we found evidence that the subcellular location of synaptic contacts of input cells onto the dendrites of starter cells does not affect spread efficiency it remains unknown what leads to the observed incomplete labeling. It is possible that this is affected by the number of synaptic contacts a presynaptic neuron makes onto the starter cell, differences in uptake receptors on the axon of distinct input cell types, or differences in postsynaptic density size (Callaway & Luo, 2015). Indeed, one limitation of our experimental design is that our light microscopy images do not allow accurate measurement of postsynaptic density size, which would require electron microscopy. Furthermore, it is important to note that we only examined spread efficiency at excitatory synapses labeled with the excitatory postsynaptic density marker PSD-95, and did not quantify the efficiency of spread to inhibitory presynaptic inputs.

Overall, we report the efficiency of spread achieved in multiple conditions consisting of distinct starter neuron types. Additionally, the described approach can be used by other researchers to test efficiency of spread across different rabies reagents, distinct input cell types, and starter neuron types not tested in this study.

## Materials and Methods

### Mouse Transgenic Lines

All experimental procedures were approved by the Salk Institute Animal Care and Use Committee. C57BL/6J mice were used as wild-type. GENSAT BAC transgenic Nr5a1-Cre (Jackson Laboratory stock # 006364), knock-in Som-IRES-Cre (Jackson Laboratory stock #013044), and knock-in Vip-IRES-Cre (Jackson Laboratory stock #028578) mice have been previously described (Harris et al., 2014; Taniguchi et al., 2011). Transgenic mice were maintained on C57BL/6J backgrounds. Mice were housed with a 12-hour light and 12-hour dark cycle and *ad libitum* access to food and water. Both male and female mice were used for experiments.

### Virus Preparation

The following AAVs were produced by the Salk GT3 Viral Core: AAV8-hSyn-FLEX-H2BmTagBFP2-oG (3.57E+13 GC/ml) and AAV8-nef-AO-66/71-TVA950 (5.25E+13 GC/mL).

AAV1-CAG.FLEX.GFPsm_myc.WPRE.SV40 (1.12E+13 GC/ml) was purchased from Addgene. EnvA+ RVdG-5PSD95eGFP-SynPhRFP (1.55E+08 IU/ml) was produced by the Salk GT3 Viral Core. AAVs and rabies virus can be purchased from the Salk Viral Vector Core.

### Animal Surgery for Virus Injection

For rabies transsynaptic spread efficiency experiments mice received AAV helper injections at postnatal day (P) 50. Mice were initially anesthetized with 2% isoflurane and maintained at 1.5% isoflurane after placement on a stereotax (David Kopf Instruments, Model 940 series) for surgery and stereotaxic injections. A small craniotomy was made with a mounted drill over the primary visual cortex of the left hemisphere using the following coordinates: 3.4 mm posterior and 2.6 mm lateral relative to bregma. For transsynaptic tracing experiments AAV8-hSyn-FLEX-H2BmTagBFP2-oG, AAV8-nef-AO-66/71-TVA950, and AAV1-CAG.FLEX.GFPsm_myc. WPRE.SV40 were mixed at a ratio of 1:40, 1:100, and 1:2 to final concentrations of 8.92E+11, 5.25E+11, and 5.6E+12 GC/ml respectively. For control experiments with no oG AAV8-nef-AO-66/71-TVA950 and AAV1-CAG.FLEX.GFPsm_myc. WPRE.SV40 were mixed to a titer matched concentration of 5.25E+11 and 5.6E+12 GC/ml. 100nl of mixture was injected into the center of V1 0.5–0.7 mm ventral from the pia using a pulled glass pipette with a tip size of 30um connected to a 1ml syringe with 18G tubing adapter and tubing. To prevent backflow, the pipette was left in the brain for 5 minutes after injection. Three weeks after AAV helper injection, 150 nl of EnvA+ RVdG-5PSD95eGFP-SynPhRFP was injected into the same site in V1. After recovery, mice were given water with ibuprofen (30mg/kg) and housed for 7 days to allow for transsynaptic rabies spread and fluorescent protein expression.

### Histology

Seven days after rabies injection, brains were harvested after transcardial perfusion using PBS followed by 4% paraformaldehyde (PFA). Brains were dissected out from skulls and post-fixed with 2% PFA and 15% sucrose in PBS at 4°C for 16–20 hours, then immersed in 30% sucrose in PBS at 4°C before sectioning. 50 um coronal brain sections were prepared using a freezing microtome. Free-floating sections were incubated at 4°C for 16 hours with rabbit anti-Myc (1:500, C3956; Sigma-Aldrich) primary antibody in PBS/0.5% normal donkey serum/0.1% Triton X-100, followed by donkey anti-rabbit conjugated to Alexa 647 (1:500, A-21206, ThermoFisher) at room temperature for 2–3 hours. Immunostained tissue sections were mounted on slides with polyvinyl alcohol mounting medium containing DABCO and allowed to air-dry overnight.

### Image Acquisition and Analysis

Individual sections were first scanned at 10x magnification using an Olympus BX63 wide-field fluorescent microscope to detect starter neurons triple positive for H2BBFP-oG, smFP_myc, and RVdG-PSD95GFP-SynPhRFP. Individual starter neurons were then imaged using a Zeiss LSM 880 Airyscan FAST Microscope with a Plan-Apochromat 63x/1.4 Oil DIC M27 objective. First, neurons were confirmed to be starter neurons by checking for expression of nuclear BFP. Z-stacks of images were acquired with a step interval of 500 nm for select dendritic domains. Dendrites immediately next to the soma and spanning 75um away were defined as proximal dendrites and representative images were acquired. Using expression of smFP_myc, dendrites were traced from the soma up to the pial surface. Dendritic regions spanning 75um from the pial surface were acquired and designated as distal domains. Images were processed and analyzed using NIH ImageJ software (FIJI). Images were first analyzed with only the far-red (smFP_myc) and green channel visible to count all PSD95GFP labeled postsynaptic densities colocalized with the far-red labeled dendritic region. After postsynaptic counts for that region were quantified, the red channel was turned on to quantify the number of presynaptic densities colocalized with postsynaptic densities. Due to the abundance of synapses across a 50um coronal section, puncta were counted manually one z-plane at a time. Wilcoxon rank-sum test for non-parametric comparisons was used for statistical analysis. For all figures: ***, p < 0.001; ns = not significant, p > 0.05.

### Statistics and reproducibility

No statistical methods were used to pre-determine sample sizes; rather sample sizes were established based upon similar studies in the literature. Stereotaxic injections were repeated in three biological replicates for each mouse line, and both male and female mice were used. All statistical tests were performed using R version 4.1.1 and details of individual tests are described in figure legends. All data are reported as the mean ± SEM.

### Data Availability

All data generated or analyzed during this study are included in the manuscript and supporting file: Source Data 1.

## ACKNOWLEDGEMENTS

We thank all Callaway Lab members for discussion. We also thank the Salk Viral Vector and Biophotonics Core staff members. This work was supported by NSF grant DBI-1707261 and NIH Diversity Supplement to R01 EY022577 (E.M.C.). M.P. was supported by the Paul and Daisy Soros Fellowship for New Americans.

## AUTHOR CONTRIBUTIONS

Conceptualization: M.P. and E.M.C.; Formal analysis: M.P.; Investigation: M.P., W.N.L., N.S.P., and P.A.M; Writing - original draft: M.P.; Writing - review and editing: M.P., W.N.L., N.S.P., P.A.M, and E.M.C.; Supervision: E.M.C.; Funding acquisition: M.P. and E.M.C.

